# Multivariate phenotypic divergence along an urbanization gradient

**DOI:** 10.1101/699017

**Authors:** James S. Santangelo, Carole Advenard, L. Ruth Rivkin, Ken A. Thompson

## Abstract

A growing body of evidence suggests that natural populations can evolve to better tolerate the novel environmental conditions associated with urban areas. Invariably, studies of adaptive divergence in urban areas examine only one or a few traits at a time from populations residing only at the most extreme urban and nonurban habitats. Thus, whether urbanization is driving divergence in many traits simultaneously in a manner that varies with the degree of urbanization remains unclear. To address this gap, we generated seed families of white clover (*Trifolium repens*) collected from 27 populations along an urbanization gradient in Toronto, Canada, and grew them up to measure multiple phenotypic traits in a common garden. Overall, urban populations had later phenology and germination, larger flowers, thinner stolons, reduced cyanogenesis, greater biomass, and were more attractive to pollinators. Pollinator observations revealed near complete turnover between urban and nonurban sites, which may explain some of the observed divergence in floral traits and phenology. Our results suggest that adaptation to urban environments involves multiple organismal traits.

## Introduction

Urbanization is rapidly changing the face of the planet. As cities develop, natural habitats experience drastic environmental changes, from increased temperatures and pollution to greater impervious surface and habitat fragmentation [1]. Evidence is accumulating to support the hypothesis that the environmental features associated with urbanization drive phenotypic differences between populations in urban and nonurban habitats [2]. For example, urban *Anolis* lizards have evolved longer limbs and more toe lamellae to improve sprint speed on the smooth artificial surfaces common in cities [3,4]. Increased impervious surfaces often lead to warmer air temperatures in cities [5], which has driven increases in thermal tolerance of urban acorn ant [6] and *Daphnia* populations [7]. These studies, and others (see table S1 in [8]) suggest that many urban populations are adapting to the novel environments created by humans.

Despite accumulating evidence of evolution in response to urban-driven environmental change, most studies focusing on phenotypic divergence associated with urbanization have examined just one or a few traits at a time [4,9–12]. However, theoretical and empirical work in other systems suggest that selection can drive the evolution of multivariate phenotypic clines along environmental gradients [13,14]. Multivariate phenotypic analyses incorporate multiple, often correlated, traits to explain overarching shifts in phenotypes across environments. Although we can make informed predictions about how particular traits might respond to urbanization from work in other systems, understanding how cities are driving phenotypic evolution requires quantifying divergence in many traits that simultaneously influence fitness. It is thus presently unclear what suites of traits are most often favored as populations become more urbanized.

Here, we investigate multivariate phenotypic evolution along an urbanization gradient in Toronto, Canada. Traits involved in plant reproduction are particularly likely to show multivariate, genetically-based phenotypic associations with urbanization between urban and nonurban populations due to the direct effect of these traits on fitness. In animal-pollinated plants, urbanization might impact fitness through changes to their pollinator communities. Both the diversity and abundance of pollinators are known to change along urbanization gradients [15,16], with positive [17–19] to negative [20] effects on pollinator visitation. Variation in pollination along urbanization gradients might drive changes in the extent of pollen limitation experienced by outcrossing plants [21,22], which can influence the strength and opportunity for pollinator-mediated selection on plant reproductive traits [23–25]. Consequently, we expect that plants could compensate for the altered pollination environment in cities by evolving altered reproductive trait values relative to non-urban plants.

## Methods

Detailed methods can be found in the online supplementary materials.

### Common garden experiment

We examined multivariate trait divergence along an urbanization gradient using white clover (*Trifolium repens*) as a model system. We grew 642 F1 generation white clover plants from seed in a common garden. These plants were distributed among 209 plant families’ from 27 populations spanning an urban-rural transect in Toronto, Ontario, Canada (Fig. 1a, [10]). The garden was located at the University of Toronto Mississauga in summer 2017. We measured several traits during our experiment, some of which are known to be under selection in this system (Table 1) [26,27].

**Table 1:** Traits (with units) measured throughout common garden experiment.

| **Trait** | **Unit** |
| --- | --- |
| **Flowing phenology** | # days to first flower |
| **Banner petal length** | mm |
| **Banner petal width** | mmm |
| **# of flowers per inflorescence** | Count |
| **# of inflorescences** | Count |
| **Defensive phenotype** | HCN+ (1) or HCN– (0) |
| **Leaf width** | mm |
| **Leaf length** | mm |
| **Stolon diameter** | mm |
| **Petiole length** | mm |
| **Peduncle length** | mm |
| **Vegetative biomass** | g |
| **Reproductive biomass** | g |
| **Relative investment in sexual reproduction** | Ratio of reproductive biomass and vegetative biomass |

**Fig. 1:**
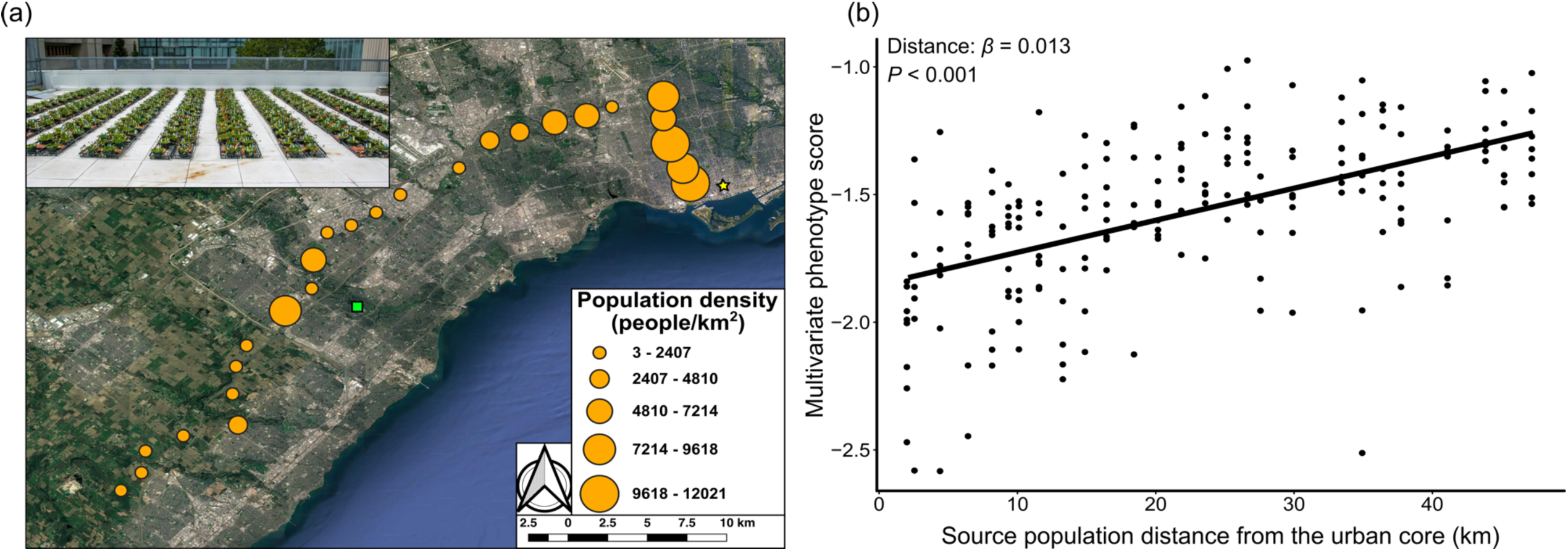
(a) 27 populations from which we collected plants (orange points). Points are scaled by population density (from [40]) within 1 km^2^. The city center (yellow star, Yonge-Dundas Square, lat: 43.6561, long: −79.3803) and common garden location (green square, lat: 43.5494, long: −79.6625) are also shown. Inset: Photograph of the common garden. (b) **cline**_**max**_ scores plotted against distance to the urban core (km).

### Field observations

We quantified variation in pollination among our study populations to identify possible mechanisms that might underlie phenotypic divergence along the urbanization gradient. We conducted pollinator observations in all study populations using two 1 m × 1 m quadrats (similar to [28]) placed in full sun and counting the number of inflorescences visited by pollinators over 20 minutes. We classified pollinators into three morphological groups (termed *morphs* hereafter): honeybees (*Apis mellifera*), bumblebees (*Bombus spp*.) and sweat bees (Halictidae).

We additionally collected twenty ripe infructescences (i.e., group of fruit) from the same populations in which we recorded pollinator observations. We counted the number of flowers and seeds in each infructescence and used the number of seeds per flower as a metric of pollen limitation (fewer seeds per flower = more pollen limitation). We generated data for the same metric from plants in the common garden. These data provide information about *in situ* variation in pollen limitation (field-collected inflorescences) and evolved differences in plants’ abilities to set seed from pollen (common garden data).

### Data analysis

We investigated whether urbanization was associated with multivariate phenotypic divergence using canonical redundancy analysis (i.e., RDA, [29]). The RDA regressed a matrix of standardized family mean trait values (209 families × 14 traits) as a response variable against a numeric vector representing the distance (in km) of populations to the urban core as the sole predictor (see below), which is correlated with % impervious surface cover (*r*_*pearson*_ = −0.63, Fig. S1a) and human population density (*r*_*pearson*_ = −0.72, Fig. S1a), has previously been shown to be correlated with cyanogenesis frequency (*r*_*pearson*_= 0.63) in the same species and region [10].

The RDA tests whether distance to the urban center explains more phenotypic variation than expected by chance. In principle, we could observe a significant RDA even if just one trait was associated with distance, but our interest lies in understanding whether multiple traits are evolving together along the transect (i.e., including traits that do not vary significantly on their own). Therefore, we used the canonical coefficients from the RDA that describe the individual contribution of phenotypic traits to the first constrained axis of the RDA (RDA1 [i.e., distance]) to calculate a multivariate phenotype score for each individual. This score, referred to as **cline**_**max**_, is the multivariate quantitative trait that shows the strongest association with distance to the urban center [13]. Regressing **cline**_**max**_ against distance to the urban center enables us to better visualize the shape of the multivariate cline and to quantify how multivariate phenotypes are changing along our urbanization gradient.

To examine variation in pollinator visitation among natural populations, we fit a linear model with the number of visits per inflorescence as a response variable, and distance, pollinator morph, and their interaction as predictors. To assess how the number of seeds per flower varied among field-collected inflorescences and plants in the common garden, we fit a model with population-mean number of seeds per flower as the response and distance, source (field-collected vs. common garden-collected), and their interaction as predictors. All data (see **Data Accessibility**) were analyzed in R version 3.6.0 [31] within the RStudio environment [32].

## Results

### Common garden

Urbanization was associated with multivariate trait divergence and the evolution of a multivariate phenotypic cline in Toronto, ON (Fig. S2 and Fig. 1b). Distance to the urban core explained 2.7% of the total variation in the multivariate phenotypic composition of populations (RDA, *F*_1, 202_ = 5.51, *P* < 0.001, Fig. S2, Fig. S3). The composite trait showing the strongest association with distance to the urban core (i.e., **cline**_**max**_) showed a positive cline along the urbanization gradient (*β* = 0.01, *t*_206_ = 9.32, *P* < 0.001, *r*^*2*^ = 0.29, Fig. 1b) that was at least as strong as any trait when analyzed individually (Fig. S4 and S5, see also standardized beta coefficients in table S1). The six traits that loaded most strongly (|loading| > 0.3, bolded in Fig. S2) onto the first axis of the RDA (RDA1)—germination, phenology, flower size, biomass, HCN, and stolon thickness—and contributed most to the composite trait showing the strongest association with distance all showed significant univariate clines in the direction predicted based on their trait loadings (Fig. S2 and Fig. S6a).

### Field observations

On average, pollinator visitation per inflorescence was greatest in urban populations (Distance: *β* = −0.032, *F*_1, 75_ = 21.96, *P* < 0.001; thick black line in Fig. 2a). Visitation rate varied with pollinator morphs (Morph: F_2, 75_ = 11.06, *P* < 0.001): bumblebees had the highest visitation rate (average = 0.63 ± 1.2 SD visits/inflorescence across all populations), followed by honey bees (average = 0.58 ± 1.05 visits/inflorescence), and sweat bees (average = 0.18 ± 0.41 visits/inflorescence). Pollinator morphs varied in their response to urbanization (Distance × Morph interaction: *F*_2, 75_ = 14.11, *P* < 0.001, Fig. 2a): bumblebee visitation rate was greatest in urban populations, whereas honeybee visitation was greatest in nonurban populations, and sweat bees showed little change in visitation across the urbanization gradient (Fig. 2a).

**Fig. 2:**
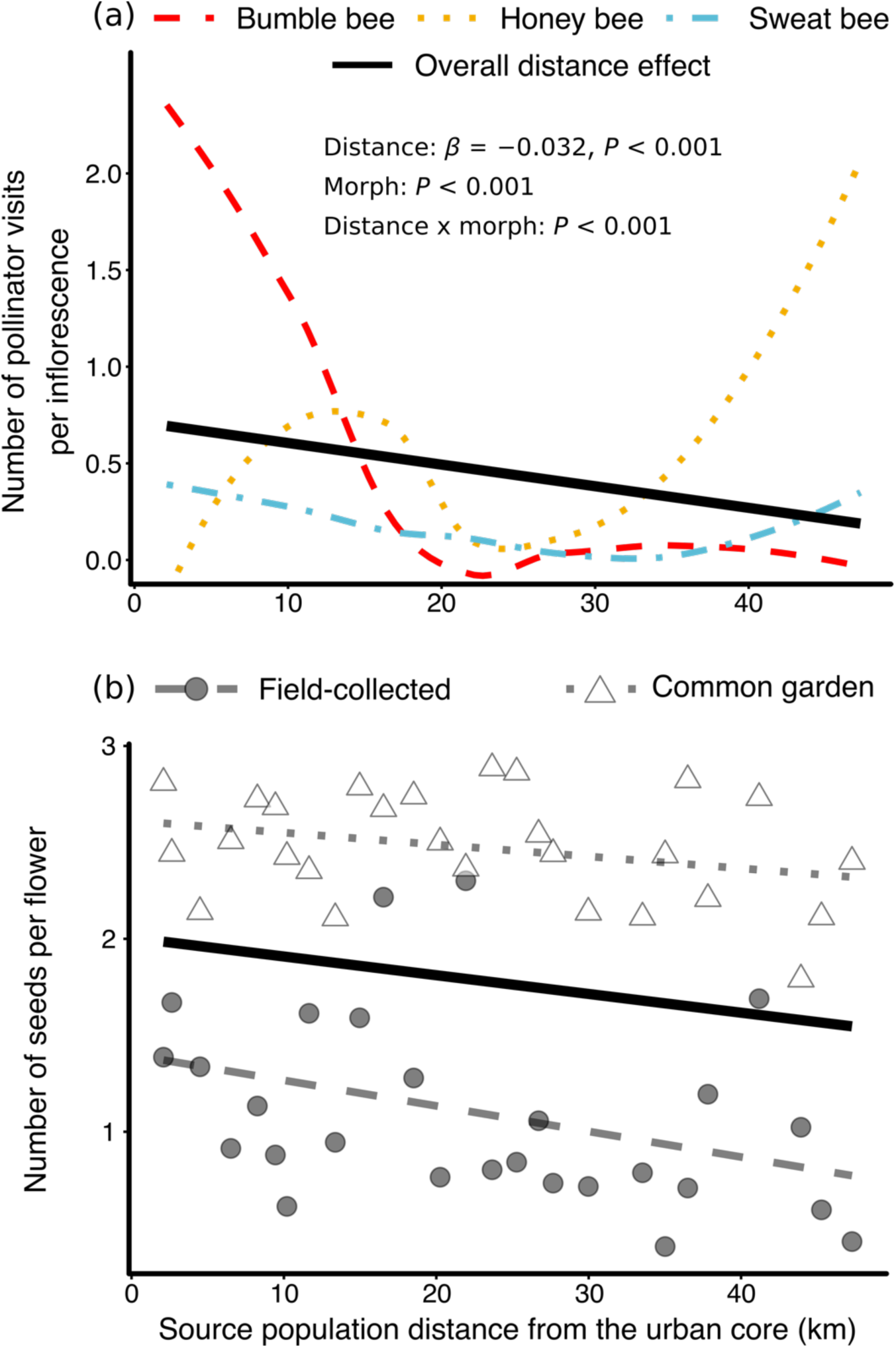
(a) Mean number of visits per inflorescence against the distance of the population from the urban core (solid black line). We also show a smoothing line (loess line) for bumblebees (red dashed), honeybees (yellow dotted), and sweat bees (blue dots/dashes) separately to illustrate the change in community composition (also see Fig. S7 for point with linear fits). (b) Number of seeds per fruit (a metric of pollen limitation) among field-collected infructescences (grey-filled circles, dashed line) and infructescences from common garden plants (white triangles, dotted line) from these same populations. The thick black line shows the overall decrease in the number of seeds per flower with increasing distance from the urban center.

While common garden infructescences were on average more thoroughly pollinated than those in natural populations (i.e., more seeds per flower; source *F*_1, 50_ = 30.04, *P* < 0.001, mean_field_ = 1.1, mean_garden_ = 2.5, Fig. 2b), the number of seeds per flower was highest in urban populations (Distance effect: *β* = −0.013, F_1, 50_ = 5.48, *P =* 0.03, Fig. 2b). Urban plants were more pollinated than non-urban plants in both the common garden and in the field (Distance × source interaction: F_1,50_ = 0.99, *P* = 0.33, Fig. 2b).

## Discussion

### Multivariate phenotypic divergence and clinal variation

Most studies of population differentiation and clinal variation along environmental gradients focus on only one or a few traits. We know much less about how suites of traits evolve in concert in response to environmental factors [13,14]. Our study shows clear phenotypic divergence and clinal variation in multivariate phenotypes along an urbanization gradient. After weighting traits based on their loadings onto RDA1 [13], distance explained 29% of the variation in the multivariate phenotype **cline**_**max**_. Urban populations had later germination and flowering (Fig. S4a and S4b), greater vegetative biomass (Fig. S4c), larger banner petals (Fig. S4d), thinner stolons (Fig. S4e), and lower HCN frequencies (Fig. S4f). Many of the traits (three of six) most strongly associated with distance to the urban core were reproductive traits, supporting the prediction that these traits are likely to show evolutionary divergence between urban and nonurban habitats, although some notable traits (e.g. reproductive biomass, number of flowers, and number of inflorescences) showed no association with urbanization.

### Urban evolution in white clover

Correlational data suggests that lower cyanogenesis in urban populations has evolved due to selection for tolerance to freezing damage [10]. Recent data suggest a cost to producing the two metabolic component of HCN (cyanogenic glucosides and linamarase) under stressful conditions (e.g., frost) [33], which may partially explain the lower HCN frequencies in urban populations. Furthermore, small-leaved plants and those with thinner stolons are more frost-tolerant [34,35], suggesting plants in urban environments should match these phenotypes if they have indeed evolved to better tolerate frost. Our data do not support the hypothesis that urban populations have evolved smaller leaves but do support urban plants having thinner stolons (Fig. S4e, table S1), offering partial support to the hypothesis that urban populations may be more frost tolerant.

### Plant evolution in urban environments

Our observed differences in flower size corroborate earlier findings of stronger directional selection for larger flowers in urban populations [36,37]. In addition, plants from urban populations were larger than those in nonurban populations, similar to urban and nonurban *Lepidium virginicum* plants grown in a common garden [12]. However, increased vegetative biomass in urban populations is not universal; in ragweed (*Ambrosia artemisiifolia*), there were no differences in plant size between urban and nonurban populations, although urban populations flowered earlier than nonurban populations [11]. This contrasts with our experiment in which urban populations flowered later than nonurban populations. These results suggest that the effects of urbanization on plant traits vary across species and there currently appears to be no particular combinations of traits consistently favored in cities.

Urban populations of white clover were less pollen limited and were visited primarily by bumblebees whereas nonurban populations were visited primarily by honeybees. A single bumblebee typically visits more white clover flowers per minute than a honeybee [38], potentially explaining the greater seed set of urban plants, and consistent with other work showing higher seed set among bumblebee-pollinated urban white clover plants [28]. The difference in pollinator community between urban and nonurban populations might explain some of the differences in floral size that have evolved if these pollinators have divergent preferences or interactions with the flowers [36,39]. In support of this hypothesis, we found that both urban common garden and field-collected plants set more seed than nonurban plants, suggesting evolved increases in seed set among urban plants which may be due to increased attractiveness to the most abundant urban pollinators. The increased pollen limitation of urban plants might have helped drive the shifts towards—or have resulted from—larger flowers in urban populations.

## Conclusion

Our results suggest that natural selection imposed by urbanization is sufficiently strong and multifarious to have driven the rapid evolution of a multivariate phenotypic cline in white clover in a large metropolitan area. The data available demonstrate variable responses of different plant taxa to urbanization, suggesting there is likely no ‘one size fits all’ solution to life in the city. Many of the traits showing divergence between urban and nonurban populations are involved in plant reproduction, and some of this divergence might be due to variation in the pollination environment. Together, our results suggest that natural selection in urban environments is rapidly refining whole-organism phenotypes to facilitate adaptation to cities.

## Acknowledgements

We thank M. Hetherington-Rauth for discussions that motivated this work. A. Dao, B. Cohan, M. Escobar, J. Viliunas, L. Miles, and S. Innes provided valuable assistance in data collection. We thank M. Kalich and B. Pitton for assistance and space in the greenhouse. Comments from the EvoEco lab at UofT Mississauga and D. Schluter’s group at UBC greatly improved the manuscript. Authors received funding from NSERC (NSERC PGS-D to JSS and NSERC CGS-D to LRR and KAT) and the Izaak Walton Killam Foundation (KAT). An NSERC Discovery Grant to Marc T. J. Johnson provided the supplies, equipment and greenhouse space rental used to carry out this project.

## Data accessibility

All code and data used throughout this manuscript are presently available for reviewers on the GitHub page for J.S.S (https://github.com/James-S-Santangelo/SIC) and will be permanently archived in a data repository following publication.

