## Supplementary material for "Multivariate phenotypic divergence along an urbanization gradient": Fig. S

### Online supplementary for: Multivariate phenotypic divergence along an urbanization gradient

**Contents:**


- Detailed methods: pp. 1 – 9
- Supplementary tables: Tables S1 (pp. 10)
- Supplementary figures: Figures S1 – S6 (pp. 11 – 16)

#### Detailed methods

*Study design*

We examined multivariate trait divergence along an urbanization gradient using white clover (*Trifolium repens*) as a model system. White clover is a perennial, herbaceous plant that is native to Eurasia but has been introduced to temperate and tropical areas globally within the past few centuries [1,2]. White clover is found in many human-altered habitats, including in cities and agricultural lands. Our focal urbanization gradient was located in Toronto, ON, Canada (population in collection year [2016] approx. 6.5 million humans). White clover is particularly suited for our research questions for several reasons. First, white clover has locally adapted for least one trait—cyanogenesis, an antiherbivore defense—in Toronto [3] and surrounding cities [4], where the frequency of cyanogenesis was positively correlated with the distance from the urban center (Fig. S2A in [3]; $R^2 = 0.40$, $P < 0.001$). Second, allele frequencies associated with local adaptation of cyanogenesis was identified as a good proxy for local adaptation generally [5], suggesting that phenotype-wide local adaptation is likely occurring in Toronto. Third, white clover possesses a considerable amount of genetic variation for many phenotypic traits [6], including investment in sexual vs. clonal reproduction [7], which can be leveraged to test predictions about the evolution of plant reproduction. Finally, white clover is typically self-incompatible, although occasional self-compatible genotypes have been documented [1,8].

*Common garden experiment*

On August 15$^{th}$, 2016, we collected plant material from 27 white clover populations spanning a 50 km gradient of urbanization intensity in Toronto (Fig. 1a). We collected up to 15 stolons (mean = 14.2) —trailing stems that give rise to foliar, floral, and root tissue—from each population for a total of 385 maternal plants, which we propagated in the glasshouse at the University of Toronto Mississauga into four-inch pots filled with Sunshine Mix Potting Soil (Sunshine mix #1; Sun Gro Horticulture, Agawam, MA, USA). Plants were kept well-watered, with ambient light, 25°C, and 50% humidity.

To minimize maternal effects and ensure that all plants in the experiment were the same age, we started our common garden using glasshouse-generated seeds. To generate these seeds, we hand-pollinated each plant with every other possible donor from its own population and harvested the seeds from these crosses when mature. Although we used individuals collected from the field as parents, this design accounts for environment-of-origin effects because nearly all of the biomass present in the seed parents was generated in the glasshouse. Throughout the article, we refer to seeds produced by an individual maternal plant within a population as a plant *family*.

To generate our common garden, we selected seeds based on their population and parent. For each of the 27 populations, we sowed five randomly selected seeds from eight randomly selected plant families ($n$ seeds sowed = 1080). Due to lack of germination and seedling mortality, we ended up with 642 plants from 209 plant families across the 27 populations (mean families per population = 7.7, mean individuals per family = 3.1). On May 15$^{th}$, 2017, we transplanted surviving seedlings into 6-inch round pots (~1.8 L) and randomized their location on the outdoor rooftop patio beside the University of Toronto Mississauga glasshouse (Fig. 1a inset). We allowed the plants to be naturally pollinated and consumed by herbivores throughout the duration of the experiment, which we ended on 30 July 2017 prior to the plants becoming root-bound in the pots.

We measured several reproductive traits during our experiment, some of which are known to be under selection in this system [6,9]. First, we recorded the number of days from germination to the production of the first open flower. Using digital calipers, we then measured the length and width of banner petals on three flowers from two inflorescences per plant. We counted the number of flowers on the first two inflorescences produced by each plant and recorded the total number of inflorescences produced over the growing season. We determined the cyanotype (i.e., cyanogenic: HCN+ or acyanogenic: HCN–) of each plant using Feigl-Anger assays [10,11].

While harvesting, we measured additional traits that might exhibit clinal variation and have diverged between urban and nonurban population. We measured the length and width of three leaves on each plant, as well as the diameter of two stolons. These traits have previously been associated with increased frost tolerance [12,13], which may be under clinal selection due to elevated frost-exposure in urban clover populations [3]. We additionally measured the length of three petioles and three peduncles to quantify plant growth form (shorter peduncles and petioles = more prostrate growth). Leaf size and petiole length were measured, relative to the leading end of a stolon, on the second most recent fully expanded leaf. The diameter of stolons was measured between the node of that leaf and the adjacent interior node. We obtained plant-level estimates of each trait by taking the mean of all measurements per individual. Finally, we harvested the sexual reproductive tissue (peduncles and inflorescences / infructescences) and aboveground vegetative tissue (everything else) separately, dried it for at least 48 h at 60 °C, and weighed it to the nearest 0.001 g. These weights were our measures of sexual and vegetative biomass.

Field observations

We quantified variation in pollination *in situ* among our study populations to identify possible mechanisms that might underlie phenotypic divergence along the urbanization gradient. This was done by conducting pollinator observations in all study populations in early August 2017. Similar to previously published methods for white clover [14], we established two 1 m × 1 m quadrats in full sun in each of the 27 populations along our urbanization gradient and counted all inflorescences with open flowers within each quadrat. We then observed each quadrat for 20 minutes and counted all pollinators and the number of inflorescences they visited during this period. We classified pollinators into three morphological groups (*hereafter*: morphs): honey bees (*Apis mellifera*), bumblebees (*Bombus spp.*) and sweat bees (Halictidae). Recent mowing in one population prevented us from collecting pollinator observations from the exact source location, so we collected data from a population within 100 m.

To complement data from pollinator observations and our field experiment, we collected twenty ripe infructescences (i.e., collection of fruit) from the same populations in which we recorded pollinator observations. Samples were collected at least 1.5 m apart. We counted the number of flowers and seeds produced by each infructescence and used the number of seeds per flower as a metric of pollen limitation (fewer seeds per flower = more pollen limitation). We generated data for the same metric from plants in the common garden by counting flowers and seeds from each of two infructescences per plant and taking the mean of these measurements as our measure of plant-level number of seeds and flowers. These data provide information about evolved differences in plants' abilities to set seed from pollen (common garden data) and *in situ* variation in pollen limitation (field-collected inflorescences).

*Data analysis*

We analyzed our data in R version 3.6.0 [15] within the RStudio environment [16] and used functions within the following packages: *tidyverse* [17], *vegan* [18], *car* [19], *lme4* [20], and *lmerTest* [21]. See **Data Accessibility** for instructions to access the data and code.

Common garden experiment

We investigated whether urbanization was associated with multivariate phenotypic divergence using canonical redundancy analysis (i.e., RDA, [22]). The RDA regressed a matrix of standardized family mean trait values (209 families × 14 traits) as a response variable against a numeric vector representing the distance of populations to the urban core as the sole predictor (see below). RDA is useful here because it allowed us to simultaneously examine divergence in multiple correlated traits (response matrix) and explore combinations of traits most strongly associated with our urbanization gradient (predictor).

Prior to the RDA, we standardized individual trait values by dividing them by their experiment-wide means and, similar to previous work examining multivariate trait divergence across environmental gradients, calculated family-mean trait values (n = 209 families) for use in our RDA [23]. Since maternal plants are independent and paternal plants were homogenized within populations, we treated families as independent in our analyses. However, results are qualitatively similar when using population means (Fig. S6). Our RDA regressed family-mean trait values (n = 14 traits) against distance to the urban center. We used distance to the urban center as a measure of urbanization because this is correlated with % impervious surface (r = –0.63, Fig. S1A) and human population density (r = –0.72, Fig. S1B), and captures appreciable variation in at least one trait (HCN) along urbanization gradients [3,4]. We calculated the distance between population and the urban center using the haversine formula [24], which calculates the distance between a pair of coordinates while accounting for Earth's curvature. We confirmed the statistical significance of distance in explaining variation of multivariate phenotypic trait divergence using 10,000 permutations of the phenotype matrix (*anova.cca* function in 'vegan', see **Results**). This approach tests the extent to which distance predicts the observed distribution of phenotypic traits relative to random rearrangements of the phenotype matrix among families.

The RDA evaluates whether distance to the urban center explains better-than-random variation in family-mean phenotypes. In principle, however, we could observe a significant RDA even if just one trait were strongly associated with distance. Thus, a significant RDA is necessary but is not alone sufficient to test our hypothesis that multivariate phenotypic divergence has occurred along our focal urbanization gradient. To probe the multivariate nature of the cline, we used the canonical coefficients from the RDA that describe the individual contribution of phenotypic traits to the first constrained axis of the RDA (RDA1 [i.e., distance]) to calculate a multivariate phenotype score for each individual. This score, referred to as **$cline_{max}$**, is the multivariate quantitative trait that shows the strongest association with distance to the urban center (Stock et al. 2014) and is calculated as:

$$cline_{max,i} = (r_1 \times Germ_i) + (r_2 \times FF_i) + (r_3 \times NumInf_i) + (r_4 \times RepBio_i)$$
$$+ (r_5 \times VegBio_i) + (r_6 \times BW_i) + (r_7 \times BL_i) + (r_8 \times PetL_i) + (r_9 \times PedL_i)$$
$$+ (r_{10} \times NumFlwr_i) + (r_{11} \times LW_i) + (r_{12} \times LL_i) + (r_{13} \times ST_i) + (r_{14} \times HCN_i)$$

Where $r_1 - r_{14}$ represent the canonical coefficients extracted from the RDA, while the remaining terms (e.g., $Germ_i$, $FF_i$, etc.) represent the 14 traits for individual *i*. As above, the calculation of **$cline_{max}$** was performed on traits standardized by dividing by the mean across all populations

[23]. Because traits are weighted by their canonical coefficients from the RDA, traits that are more strongly associated with distance to the urban core contribute more strongly to a population's $\mathbf{cline_{max}}$ score. Regressing $\mathbf{cline_{max}}$ against distance to the urban center thus enables us to better visualize the shape of the multivariate cline and better quantify how multivariate phenotypes are changing along our urbanization gradient.

To complement the multivariate approach taken above, we used univariate models to examine trait change along our urbanization gradient. This allowed us to determine if the traits that contributed most strongly to multivariate trait divergence and clinal variation also showed strong clines when analyzed individually. In these models, we regressed family-mean trait values against distance to urban core as the sole predictor. Second, we used the univariate model to test the specific prediction that urbanization alters relative investment in sexual vs. clonal reproduction. In this model, we used the ratio of sexual biomass to vegetative biomass as our response variable and distance to the urban center as our predictor. Plants with a higher ratio invest more strongly into vegetative biomass and a value of zero means that a plant produced no sexual biomass.

<u>Field observations</u>

We used linear models to assess how pollinator visitation and the number of seeds per flower among field-collected and common garden plants varied along the urbanization gradient. To examine variation in pollinator visitation, we fit a linear model with the number of visits per inflorescence as a response variable, and distance, pollinator morph, and their interaction as fixed effect predictors. We obtained parameter estimates and *P*-values using type III sums of squares. In this model, a significant effect of distance suggests that pollinator visitations vary along our urbanization gradient, a significant effect of morph suggests that pollinator taxa vary in their overall visitation rates, while a significant distance × morph interaction suggests that visitation rates by at least two pollinator taxa change in different ways along the urbanization gradient.

To assess how the number of seeds per flower varied among field-collected inflorescences and plants in the common garden, we fit a model with population-mean number of seeds per flower as the response and distance, source (field-collected vs. common garden-collected), and their interaction as fixed-effect predictors. We obtained parameter estimates and *P*-values using type III sums of squares. In this model, a significant effect of distance suggests that the number of seeds per flowers varies with urbanization, a significant effect of source suggests that field-collected and common garden plants vary in the number of seeds per flower, while a significant distance × source interaction suggests that the effects of population origin on pollen limitation is different in the common garden than at the field sites. This interaction term provides insight into whether variation in pollen limitation was due to extrinsic (reduced pollination) or intrinsic (e.g., increased attractiveness to pollinators) factors.

### Tables

**Table S1:** Mean of family-mean trait values for 14 traits included in the multivariate analysis of differentiation across our urbanization gradient. Also shown are the beta coefficients, *P*-values, and $R^2$ values from models with unstandardized or mean-standardized traits as the response variable and distance of source population to the urban center as the sole predictor. The figure column references the figure showing the univariate cline for each trait.

| Trait | Mean | Unstandardized $\beta$ | Standardized $\beta$ | *P*-value | $R^2$ | Figure |
|---|---|---|---|---|---|---|
| **Days to germination** | 4.96 | -0.055 | -0.0112 | < 0.001 | 0.1583 | S4a |
| **Days to first flower** | 62.35 | -0.067 | -0.0011 | 0.0153 | 0.0281 | S4b |
| **Vegetative biomass (g)** | 3.65 | -0.0196 | -0.0054 | < 0.001 | 0.0653 | S4c |
| **Banner petal length (mm)** | 6.19 | -0.0048 | -0.0008 | 0.0463 | 0.0191 | S4d |
| **Stolon thickness (mm)** | 1.38 | 0.0021 | 0.0015 | 0.0047 | 0.038 | S4e |
| **Frequency of HCN** | 0.25 | 0.004 | NA* | 0.0122 | 0.0307 | S4f |
| **Number of inflorescences** | 20.89 | -0.0023 | -0.0001 | 0.9606 | 0 | S5a |
| **Reproductive biomass (g)** | 1.61 | -0.0004 | -0.0003 | 0.8942 | 0.0001 | S5b |
| **Banner petal width (mm)** | 3.56 | -0.0021 | -0.0006 | 0.0884 | 0.014 | S5c |
| **Peduncle length (mm)** | 101.98 | 0.0694 | 0.0007 | 0.4183 | 0.0032 | S5d |
| **Number of flowers** | 46.17 | 0.0322 | 0.0007 | 0.5659 | 0.0016 | S5e |
| **Leaf width (mm)** | 11.02 | -0.0012 | -0.0001 | 0.8617 | 0.0001 | S5f |
| **Leaf length (mm)** | 12.59 | -0.0009 | -0.0001 | 0.8965 | 0.0001 | S5g |
| **Petiole length (mm)** | 446.76 | 0.3686 | 0.0008 | 0.4322 | 0.003 | S5h |
| **$Cline_{Max}$** | NA | NA | 0.013 | < 0.001 | 0.30 | 1b |

* HCN frequency was not mean standardized since it was encoded as binary (HCN+ = 1; HCN− = 0)

### Figures

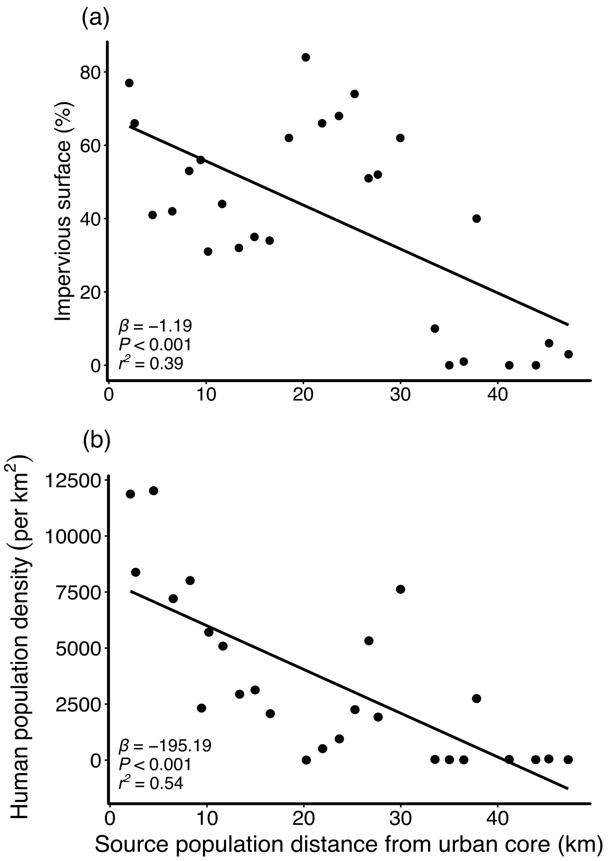


**Fig. S1:** Regressions of percent impervious surface (a) and human population density (b) with distance to the urban core. Each point represents one of 27 populations. Human population density within 1 $km^2$ were obtained from [25] and percent impervious surface within 1 $km^2$ were obtained from [26].

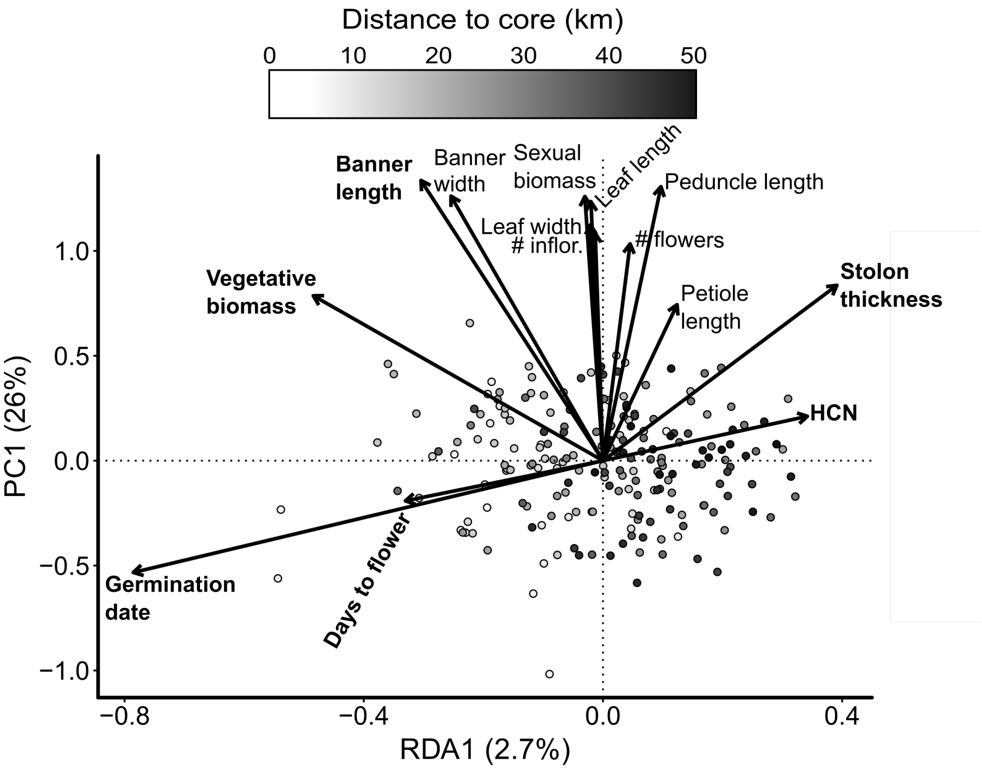


**Fig. S2:** RDA triplot displaying the position of each family along the constrained axis (RDA1) and first unconstrained axis (PC1) of the canonical redundancy analysis. Families are shaded based on their distance to the urban center. Solid arrows represent standardized trait loadings: the direction of arrows relates to the orientation of axes with which they are most strongly correlated while the length of the arrow corresponds to the strength of that association. Bolded traits are significant ($P < 0.05$) when analyzed as univariate clines (Fig. S4). Populations further from the city center (i.e. right along RDA1) had higher HCN frequencies, smaller banner petal widths and lengths, lower vegetative biomass, flowered earlier, and germinated earlier. Other traits were not strongly associated with distance. Note that average seeds per flower (Fig. 2b) was not included in the RDA due to too much missing data at the family level.

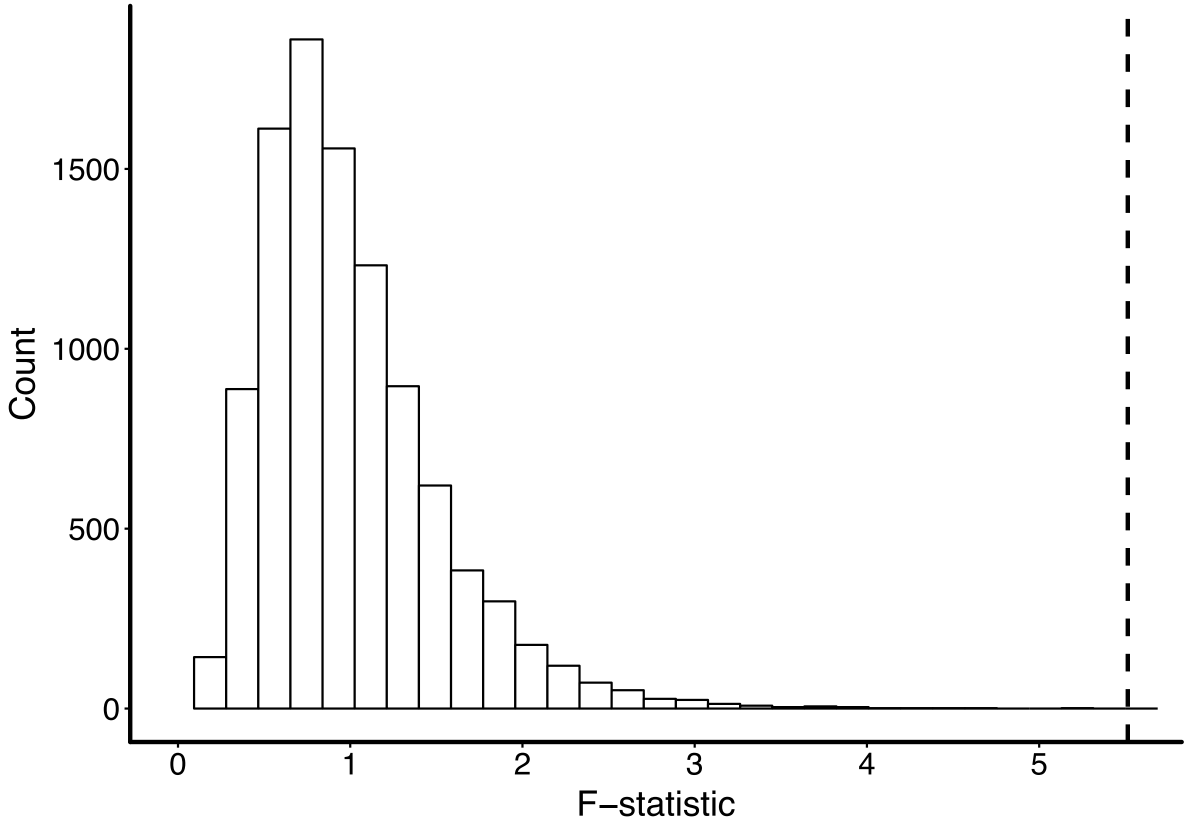


**Fig. S3:** Distribution of 10,000 simulated *F*-statistics following row-wise permutations of the RDA trait (i.e., response) matrix. The vertical dashed line shows the observed F-statistic. Simulated and observed values were obtained using the *anova.cca* function from the *vegan* R package.

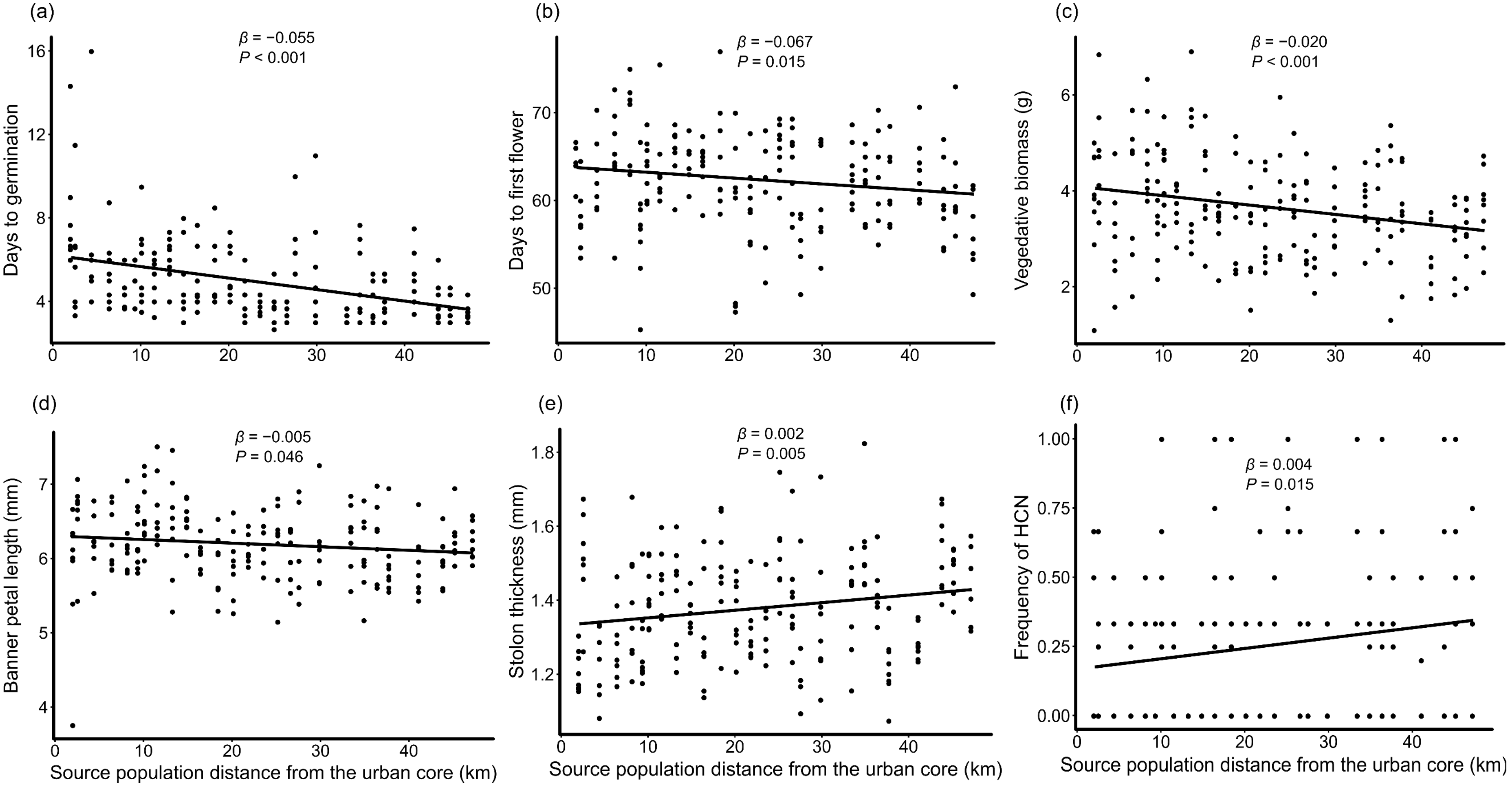


**Fig. S4:** Significant ($P < 0.05$) univariate regressions of family-mean trait values against source population distance from the urban core. Each point represents the mean trait value across individuals in the same plant family and population.

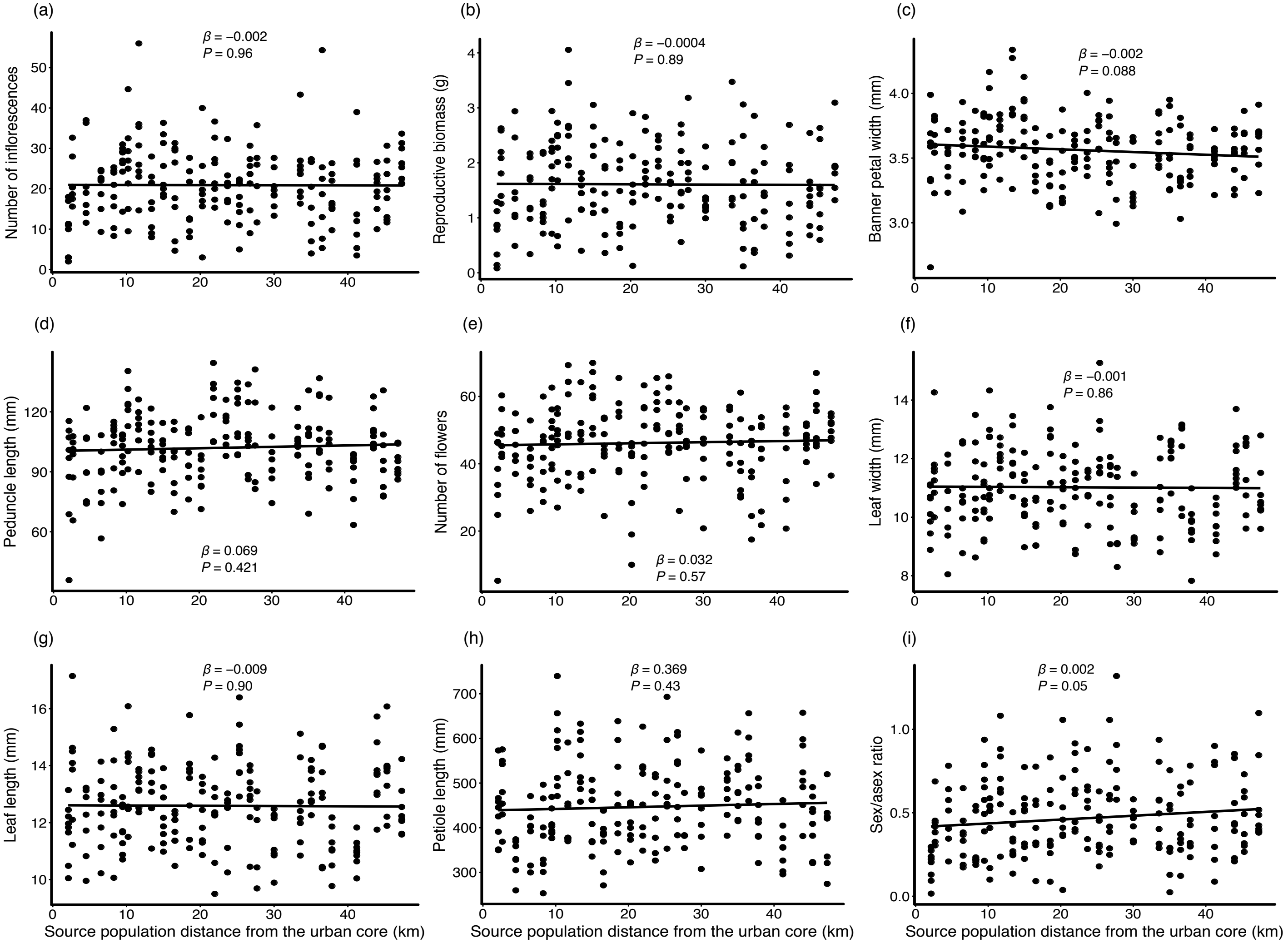


**Fig. S5:** Non-significant ($P > 0.05$) univariate regressions of family-mean trait values against source population distance from the urban core. Each point represents the mean trait value across individuals in the same plant family and population. Note that sex/asex ratio (i) was not included in the RDA since it is a linear combination of traits already included in the analysis.

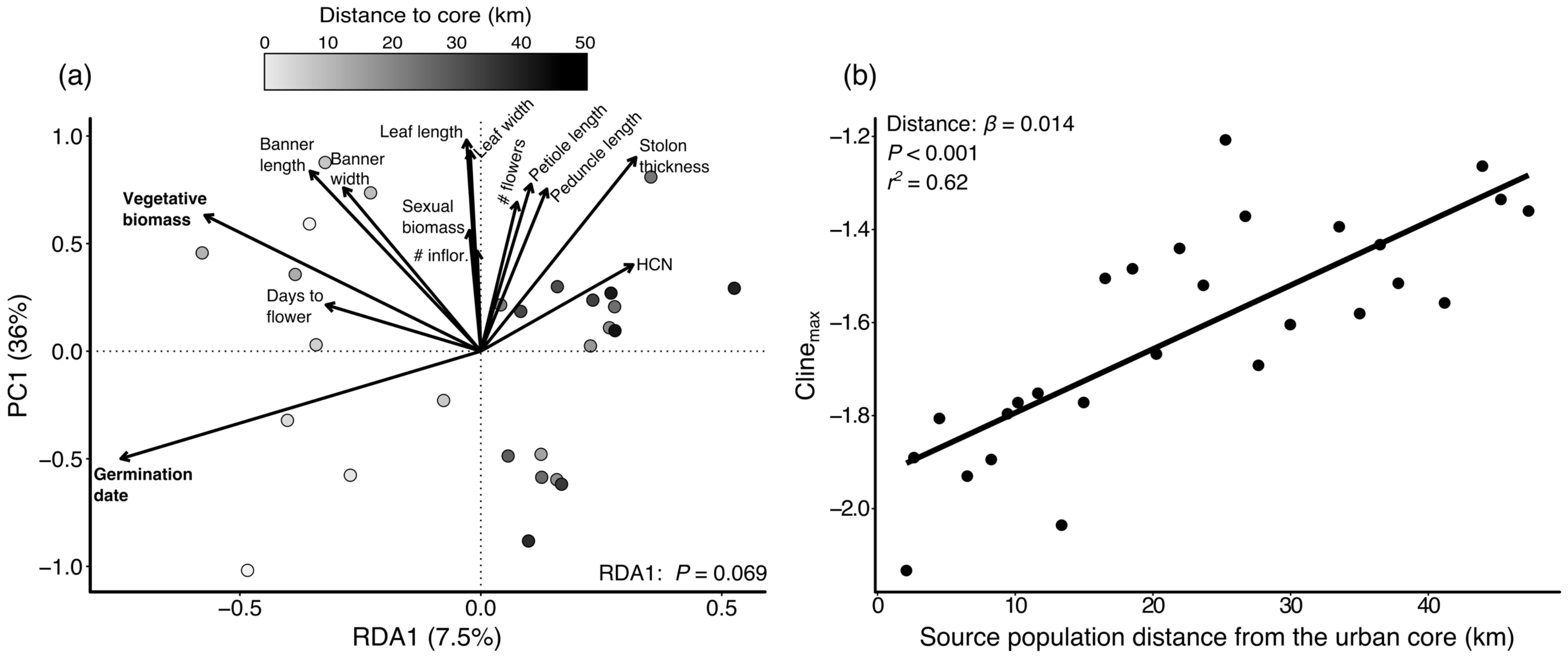


**Fig. S6: Multivariate phenotypic divergence along an urbanization gradient using population means.** Panel (a) shows an RDA triplot displaying the position of each population along the constrained axis (RDA1) and first unconstrained axis (PC1) of the canonical redundancy analysis. Populations are shaded based on their distance to the urban center. Solid arrows represent standardized trait loadings: the direction of arrows relates to the orientation of axes with which they are most strongly correlated while the length of the arrow corresponds to the strength of that association. Bolded traits are significant ($P < 0.05$) when analyzed as univariate clines. Populations further from the city center (i.e. right along RDA1) had faster germination and produced less vegetative biomass. Other traits were not strongly associated with distance. Panel (b) shows the population **$\text{cline}_{\text{max}}$** scores plotted against distance to the urban core (km). The distance to the city center was a significant prediction of **$\text{cline}_{\text{max}}$,** the multivariate trait showing the strongest association with distance based on RDA scores.

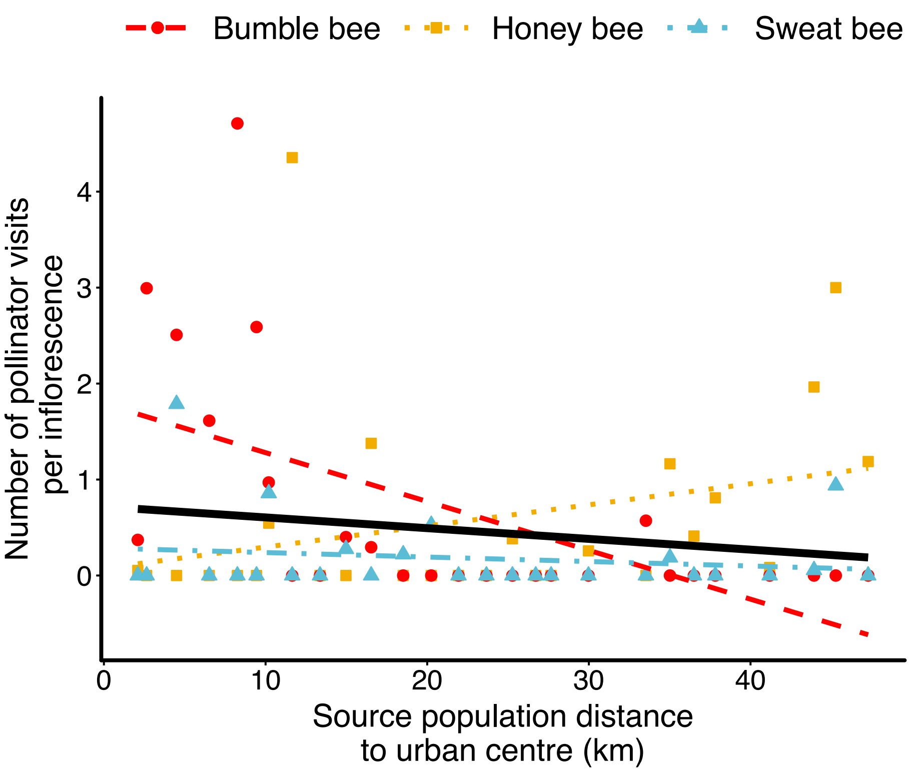


**Fig. S7:** Relationship between the mean number of visits per inflorescence and the distance of the population from the urban core (solid black line). Because the effects of distance on the number of pollinator visits varied across pollinator taxa, we also show a separate linear fit for bumblebees (red dashed line with points), honeybees (yellow dotted line with squares), and sweat bees (blue dots/dashes with triangles) to illustrate the change in community composition.
